# Stomatal patterning is shaped by the interplay with giant cell patterning in Arabidopsis

**DOI:** 10.64898/2026.04.30.721859

**Authors:** Gauthier Weissbart, Frances K. Clark, Adrienne H. K. Roeder, Pau Formosa-Jordan

**Author notes:** Laboratoire Reproduction et Développement des Plantes (RDP), Université de Lyon, ENS de Lyon, UCB Lyon 1, CNRS, INRAE, Inria, 69342, Lyon, France.

## Abstract

In developing tissues, cells differentiate into distinct cell types and form complex spatial patterns. How distinct patterning systems interact during tissue growth to shape tissue composition and spatial organization remains poorly understood. Here, we investigate this question in the abaxial leaf epidermis of *Arabidopsis thaliana*, in which the same pool of progenitor cells gives rise to stomata, pavement cells, and giant cells. Using a quantitative approach combining Euclidean and network-based spatial analysis, we show that stomatal number and density are robust to reduced endoreduplication, whereas forced endoreduplication actively competes with the stomatal lineage to reduce stomatal number. Furthermore, we show that the stomatal spatial pattern is also shaped by the broader tissue context such as cell growth, cell division, and giant cell patterning, with distinct consequences for stomatal spatial distribution and cellular arrangement. Our results highlight that the interplay between patterning systems must be considered to understand how tissue organization is established.

## Introduction

During development, initially identical cells differentiate into distinct cell types and form spatial patterns. Multiple studies have investigated how one cell type becomes specified from progenitor cells. However, tissues are often composed of more than two cell types, each of which can originate from a different signalling pathway and undergo its own patterning process. How the underlying patterning processess leading to each cell type interact to shape tissue composition and spatial organization remains unclear. The abaxial leaf epidermis of *Arabidopsis thaliana* provides an ideal model to address this question. It is composed of three main cell types — stomatal guard cells [1, 2], which appear in pairs to form the stomatal complexes, pavement cells, and giant cells [3, 4, 5] — each with distinct functions [6]. Stomatal complexes, herein referred to as stomata, form a relatively regular spatial pattern, separated by at least one cell due to lateral inhibition and lineage-specific division orientation [7, 8, 9]. Their patterns have been largely investigated in relation to their function related to gas exchange and water use efficiency [10, 11, 12]. Giant cells are large endoreduplicated cells scattered throughout the tissue [3, 4], which become clustered in the mature epidermis as a consequence of surrounding cell divisions [5, 13]. Although stomatal patterning and giant cell patterning have been studied independently, whether and how these two patterning systems interact during tissue growth to shape tissue organization remains an open question.

All cell types in the leaf epidermis arise from an initial pool of undifferentiated protodermal cells. Although it is generally acknowledged that a gradient of cell proliferation is followed by cell differentiation and the expansion of pavement cells from the tip to the base of the leaf [14, 6], these processes can occur concurrently and simultaneously, and are coordinated in a complex manner [15, 16, 17]. A subset of cells accumulate SPCH protein and enters the stomatal lineage, a multi-stage program of asymmetric divisions generating meristemoids and stomatal lineage ground cells (SLGCs), ultimately producing both stomata and pavement cells [8, 7]. Cells from the stomatal lineage account for approximately half of the epidermal cells [18]. The other protodermal cells can enter the endoreduplication cycle through the *ATML1* pathway, forming giant cells [3, 4, 5], while other becomes pavement cells. These two programs, stomatal entry and giant cell specification, might therefore compete for the same initial pool of protodermal cells. Yet, whether giant cell patterning influences stomatal number or spatial organization, either through direct competition or by altering the stomatal lineage program, remains unclear.

The spatial organization of stomata has been investigated in several studies [18, 19, 9], but almost exclusively from a point-pattern perspective measuring Euclidean distances [20, 21]. This approach cannot distinguish whether observed spatial differences reflect a discrete patterning mechanism or is the consequence of cell growth and shape changes. Moreover, how cell division and endoreduplication influence stomatal spatial patterning has not been extensively investigated.

Here, we adopt a quantitative image analysis approach combining Euclidean and network-based spatial analysis to understand how giant cell patterning and cell division influence stomatal number and spatial organization. We apply this approach to segmented images of leaf epidermal tissues in wild-type and mutant genotypes of the giant cell specification pathway [5]. We show that in addition to the stomatal signaling pathway, the stomatal spatial pattern is also shaped by the broader tissue context: cell growth, cell division, and giant cell patterning. These results highlight that the interplay between patterning systems must be considered to fully understand how is the mixture of cell types achieved in leaves and how is stomatal organization established.

## Results

### Stomatal number and density are robust to reduced endoreduplication and increased cell division

It has been shown that a decrease in endoreduplication is compensated by the presence of more small cells ([4, 5]. However, it is unclear whether these small cells are pavement cells formed after mitotic divisions or stomatal lineage cells formed through new stomatal lineage entries. Here, we quantified the cell type composition in published images of leaf regions of mutants affecting the giant cell specification pathway (Fig. 1A) ([5]. We quantified the fraction, the absolute number (estimated using leaf size measurements [5]), and the density (number divided by section area) of each cell types across three replicates.

**Figure 1:**
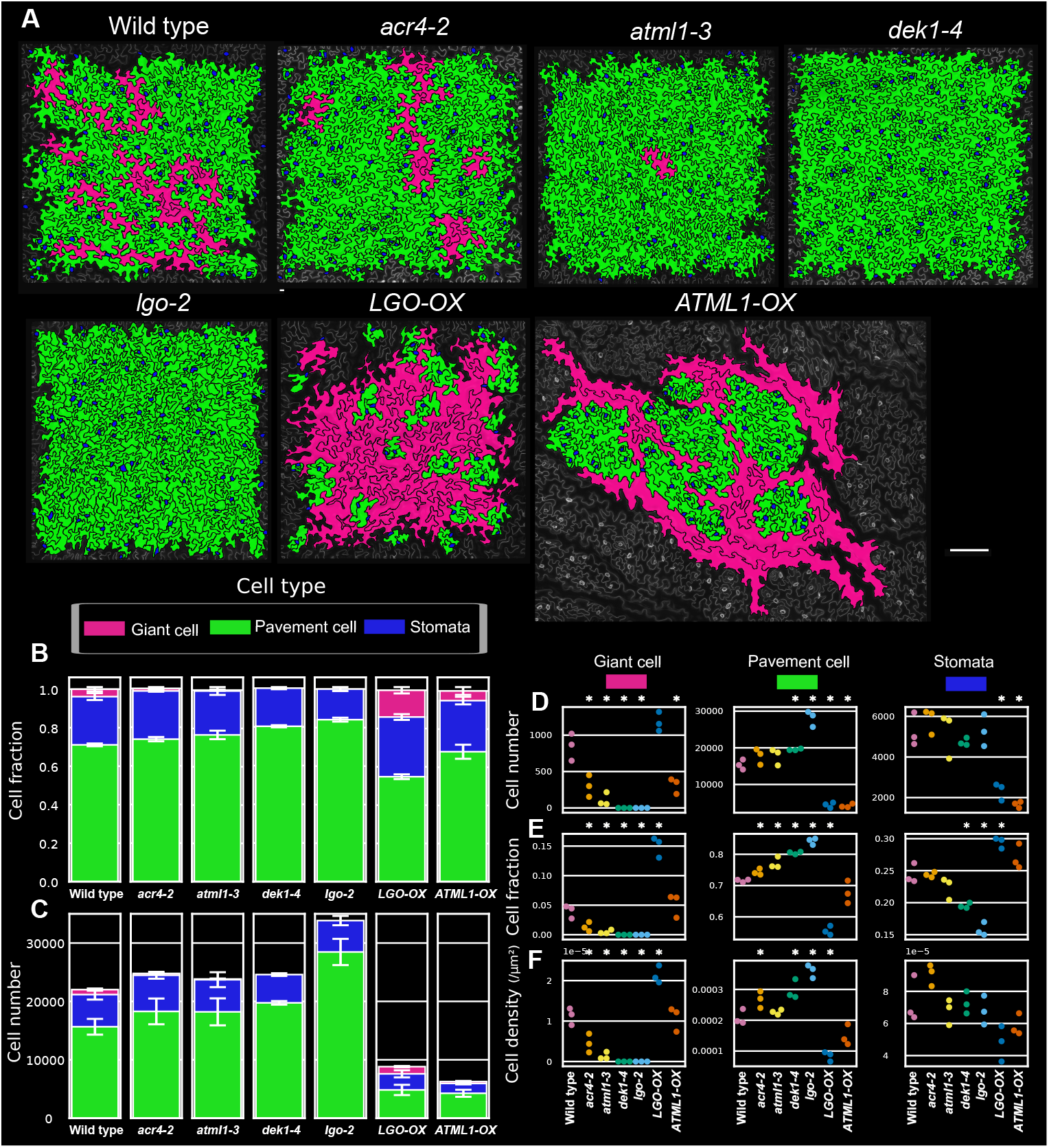
Giant cell patterning alters the cell type composition in the leaf epidermis. (A) Segmented regions of the abaxial side of leaf epidermal tissues at 25 dpg, after cell type classification, in the genotypes of the giant cell specification pathway: wild type, *acr4-2, atml1-3, dek1-4, lgo-2, LGO-OX*, and *ATML1-OX*. These panels are reproduced from [5]. (B) Cell fraction (or proportion) for each cell type in each genotype. The mean was averaged over three individual replicates. (C) Estimated cell number in the entire leaf blade for each cell type in each genotype. To estimate these numbers, the number of cells in the segmented tissues was multiplied by the leaf size and a factor of 0.9 (estimation of the blade surface) divided by the area of the segmented region. The mean was averaged over three individual replicates. (D) Estimated cell numbers in the entire leaf blade in each replicate for giant cells, pavement cells and stomata. (E) Cell fractions in each replicate for giant cells, pavement cells and stomata. (F) Cell density in each replicate for giant cells, pavement cells and stomata. In (A–C), giant cells are labeled in magenta, pavement cells in green, and stomata in blue. In (D–F), Statistical tests were performed using t-tests. * represents a significance with a p-value < 0.05.

In loss-of-function mutants, in which the giant cell fraction is significantly reduced, the total number of cells was, as expected, increased, especially in *lgo-2*. The cell type composition was altered, with a significant increase in the fraction of pavement cells (Fig. 1B, E) in all mutants, and a decrease in the stomatal index (i.e., the fraction of stomata cells in the tissue) in *dek1-4* and *lgo-2*. We observe that reducing giant cell numbers increase the pavement cell fraction and decrease the stomatal fraction. In this case, a reduced stomatal fraction is expected regardless of whether a fate competition exists between stomatal lineage entry and giant cell fate, since stomatal lineage initiation typically produces more pavement cells than stomata (about 1.6 pavement cells per stomata), leading to a decrease in stomatal fraction. Surprisingly, however, the absolute number of stomata remained approximately constant in all loss-of-function studied mutants (Fig. 1D). Similarly, while pavement cell density increased, stomatal density remained largely unchanged. Therefore, a reduction in giant cell formation is mostly compensated by the production of additional pavement cells, but not by an increase in stomatal number. This result challenges the possibility that that giant cell specification and stomatal initiation compete for a finite pool of protodermal cells, and is consistent with previous findings in *lgo-2* [22]. The absolute number of pavement cells (Fig. 1C, D) significantly increased in all loss-of-function mutants. In *lgo-2*, we observed a compensation of approximately 16 additional pavement cells for each reduction of one giant cell compared to wild type.

Taken together, these results show that reducing the giant cell fraction alters tissue composition primarily through increased pavement cell production. This indicates that stomatal number is robust to a decrease in giant cells and increased cell division, suggesting that the initiation of stomatal patterning might not depend on the discrete number of cells and be regulated at the continuous space (i.e., in micron or microns square units instead of cell units) or at the tissue level if both patterning systems occur simultaneously. However, these results should be interpreted with caution, as the reduction in giant cell number in these mutants is relatively small, and subtle changes in stomatal number may not be statistically detectable.

### Increasing giant cell number reduces both stomatal and pavement cell numbers

We then investigated how promoting giant cell fate through the overexpression of ATML1 (*ATML1-OX*) and LGO (*LGO-OX*) impacts tissue cell type composition. In *ATML1-OX* and *LGO-OX* leaves, the total number of cells was strikingly reduced, with an estimate of 2–3 times fewer cells compared to wild type. In contrast to loss-of-function mutants, we observed a significant reduction in the absolute number of stomata (Fig. 1D). As a result, the stomatal fraction was increased, whereas the pavement cell fraction was reduced (Fig. 1E). This suggests that, when giant cell commitment is enhanced, some cells adopt a giant cell fate instead of entering the stomatal lineage, raising the possibility of competition for cell fate.

Interestingly, the two overexpression lines show distinct effects on tissue composition. In *ATML1-OX*, giant cell fraction and overall cell type composition remains relatively similar to wild type. By contrast, *LGO-OX* exhibits a much higher proportion of giant cells and giant cell density, suggesting that additional cells are recruited into the giant cell fate. We wondered whether stomatal lineage cells can also become giant cells. In wild type, some stomatal lineage cells enlarge but do not reach the size threshold to be classified as giant cells. In *LGO-OX*, we observed several stomata surrounded only by giant cells, indicating that some stomatal lineage cells can endoreduplicate and become giant. This may reduce stomatal production, as cells that endoreduplicate limit lineage amplification in the stomatal lineage program. By contrast, *ATML1-OX* did not show obvious giant cells surrounding stomata. Instead, pavement cells appeared smaller and were likely less impacted by endoreduplication. This suggests that *ATML1-OX* does not promote giant cell fate within the stomatal lineage.

Overall, this quantification indicates that giant cell formation can lead to a reduction of stomatal production, either by reducing entry into the stomatal lineage or potentially by altering its stomatal lineage progression in *LGO-OX*. Therefore, these results support the existence of interplay between stomatal and giant cell patterning processes. However, alternative explanations, such as earlier termination of organ growth or regulation at the organ level, could also contribute to the reduction in cell number.

### Distinct stomatal spatial distributions mask a similar cellular arrangement due to cell size and shape

We have shown that giant cell formation via endoreduplication plays a role in the mixture of cell types in the epidermis, by influencing the proportions and numbers of stomata and pavement cells. Because we observed some differences in the stomatal spatial pattern in our phenotypes (Fig. 1A), we asked how giant cell patterning influences the spatial patterning of stomata.

To investigate the impact of giant cell patterning on the stomatal spatial pattern, spatial quantification are required. Previous studies quantified stomatal patterns in Euclidean space and concluded that stomata are locally ordered but randomly distributed at larger scales [19, 18, 20]. However these analyses treated cells as points, potentially masking order at the cellular scale due to cell size and shape heterogeneity [23]. Here, we quantify stomatal patterns in both Euclidean space (referred to as “spatial distribution”) and network space (referred to as “cellular arrangement”) to be able to distinguish changes in patterning at the cellular level from effects of cell size and shape.

To investigate the effect of cell size and shape on stomatal distribution, we first compared the stomatal pattern in leaves and sepals, which exhibit very distinct cell shapes despite comparable cell type compositions. (Fig. 2A). In the Euclidean space, the average nearest neighbor (ANN) between stomata was shorter in sepals (ANN 40 µm) compared with mature leaves (ANN=65 µm) (Fig. 2B), with less spread NN and k-nearest neighbor distance distributions, indicating that stomata are spatially more packed in sepals (Fig. 2B, C), and that fewer stomata are at a large distance from other. At larger scales, stomatal patterns were randomly organized in both organs, but randomness appeared at a shorter distance in sepalsn as indicated by the pair correlation functions (Fig. 2D) (g close to 1 for r > 100 µm in leaves and for r > 30 µm in sepals).

**Figure 2:**
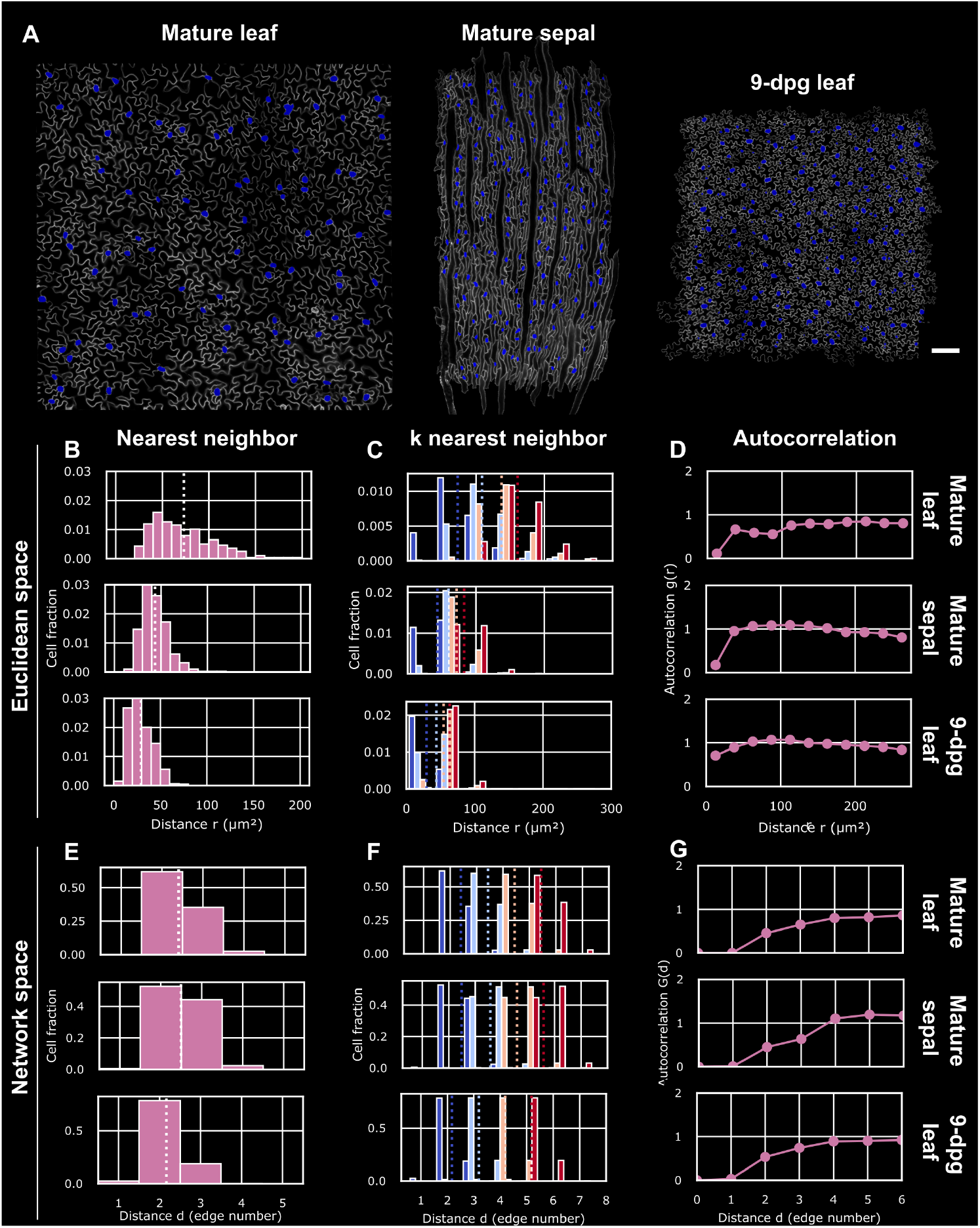
Distinct stomatal spatial distributions in Euclidean space mask a similar cellular arrangement in network space. (A) Segmented regions of a mature leaf (left), a mature sepal (middle), and a 9-dpg leaf (right) (stomata labeled in blue). (B–D) Spatial measures in the Euclidean space to quantify the spatial distribution of stomata in the mature leaf (top), the mature sepal (middle), and the 9-dpg leaf (bottom). Stomata are considered here as point coordinates in the Euclidean space (see wild type in Fig. 3A). (B) Nearest neighbor (NN) distributions (in µm) of stomata. The vertical dotted line indicates the average nearest neighbor (ANN). (C) K nearest neighbor distributions of stomata with k from 1 (first nearest neighbor) to 4 (fourth nearest neighbor): k = 1 is dark blue, k = 2 is light blue, k = 3 is light red, and k = 4 is dark red. Vertical dotted lines indicate each k average nearest neighbor (kANN). (D) Pair correlation function (see Methods) (g < 1 indicates order, whereas g > 1 indicates clustering). (E–G) Spatial measures in the network space to quantify the stomata spatial arrangement in the mature leaf (top), the mature sepal (middle), and the 9-dpg leaf (bottom). Cells are here represented within the Network space (see wild type in Fig. 4A). (E–G) are analogous measures of (B–D) in the network space. See also Figs. 3, 4 and Figs. S2 and 5.

These findings highlight a difference in the spatial distribution of stomata between these organs. However, this might be attributed to larger and more isotropic pavement cells in the mature leaf, which may contribute to increasing the distance between stomata. To investigate this hypothesis, we analyzed the stomatal pattern in the developing leaf at 9 dpg, where pavement cells are less expanded but stomatal differentiation is nearly complete. Stomata were indeed closer spatially at 9 dpg (ANN = 35 µm), and their overall spatial distribution was strikingly similar to those observed in sepals (Fig. 2B, D). This suggests that the expansion of pavement cells, including giant cells, is responsible for spacing stomata in the Euclidean space as the leaf grows.

However, because the observed differences might also be attributable to a different topological arrangement within the tissue, we analyzed the analogous observables in network space (Fig. 2E–G). We found that the ANN and the k-NN distributions were relatively similar in across organs. The estimated pair correlation function adapted to network space (Fig. 2G) (see Methods) showed randomness after approximately 3 cells in both organs. A notable difference remained: in sepals, nearest stomata were more frequently found at a distance of 2 cells (Fig. 2E–F), suggesting that sepal SLGCs undergo fewer spacing divisions. Despite this, the mature leaf, the mature sepals, and the developing leaf exhibited more similarities in network space than in Euclidean space (Fig. 2B–G). This supports the idea that the same patterning mechanisms occur in leaves and sepals. These results demonstrate the complementary value of the two spatial perspectives. The differences in stomatal spatial distribution in Euclidean space between mature leaves and sepals, and between 9 dpg and 25 dpg leaves, are mostly due to differences in cell sizes and shapes. Cell expansion plays a role in spatially distributing stomata through a physical size effect, independently of the stomatal patterning mechanism itself.

### Loss of giant cells alters stomatal arrangement but preserves stomatal distribution

Because stomatal entry and giant cell specification can occur simultaneously during tissue growth, we asked whether reducing giant cell number affects stomatal spatial organization. We quantified the stomatal pattern in loss-of-function mutants using both Euclidean and network metrics (Figs. 3, 4 and Fig. S2).

**Figure 3:**
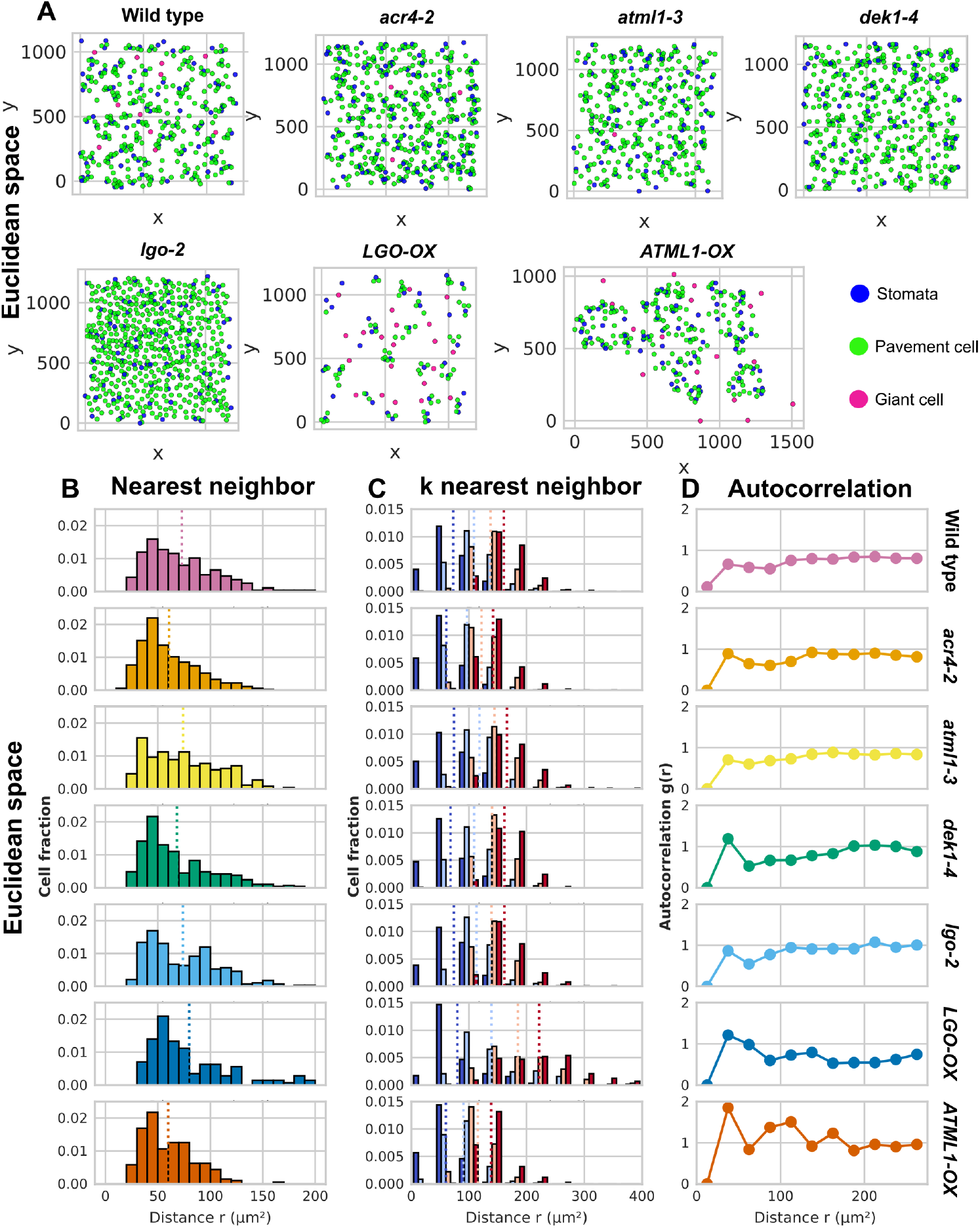
Stomatal spatial distribution in the Euclidean space is unchanged in giant cell loss-of-function mutants but altered when overexpressing *ATML1* and *LGO*. (A) Cellular patterns as point patterns. Scatter plots of cell center coordinates of one replicate for each genotype in the Euclidean space. (B–D) Spatial measures in the Euclidean space to quantify the spatial distribution (in µm) of stomata in: wild type, *acr4-2, atml1-3, dek1-4, lgo-2, LGO-OX*, and *ATML1-OX*. (B) Nearest neighbor (NN) distributions of stomata. The vertical dotted line indicates the average nearest neighbor (ANN). (C) K nearest neighbor distributions of stomata with k from 1 (first nearest neighbor) to 4 (fourth nearest neighbor): k = 1 (first nearest neighbor) is dark blue, k = 2 is light blue, k = 3 is light red, and k = 4 is dark red. Vertical dotted lines indicate the k average nearest neighbor (kANN). (D) Pair correlation function computed as the average number of cells found between r and r + dr in comparison to a random point pattern (g > 1 indicates clustering, g < 1 indicates dispersion, and g ≈ 1 indicates randomness).

**Figure 4:**
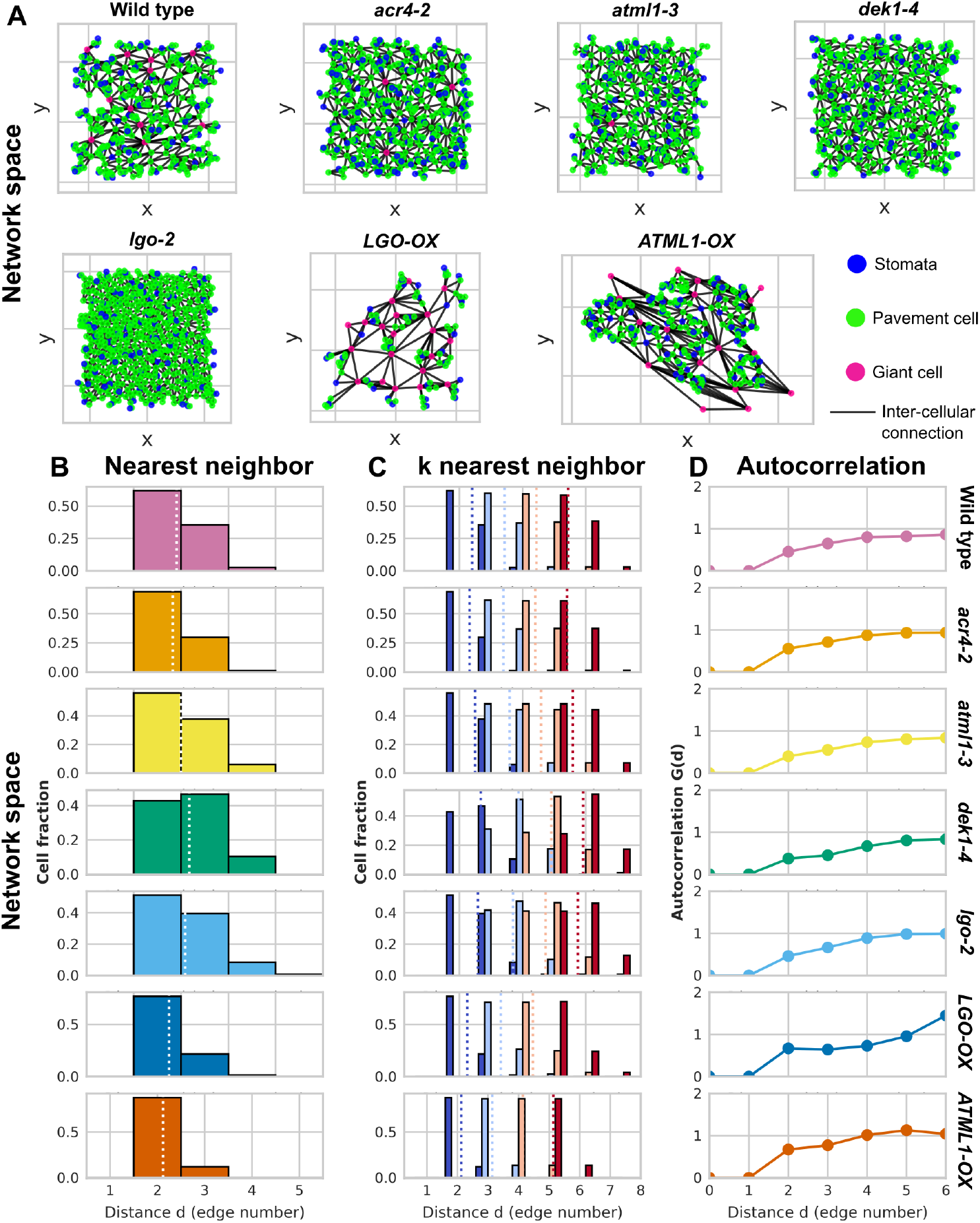
Stomatal spatial arrangement in the network space is influenced by giant cell patterning. (A) Cellular patterns as networks. Graph of one replicate for each genotype, with points representing cell center coordinates and edges representing cell connections in the cellular network. (B–D) Spatial measures in the network space to quantify the spatial arrangement (in edge number) of stomata in: wild type, *acr4-2, atml1-3, dek1-4, lgo-2, LGO-OX*, and *ATML1-OX*. (B) Nearest neighbor (NN) distributions of stomata. The vertical dotted line indicates the average nearest neighbor (ANN). (C) K nearest neighbor distributions of stomata with k from 1 (first nearest neighbor) to 4 (fourth nearest neighbor): k = 1 (first nearest neighbor) is dark blue, k = 2 is light blue, k = 3 is light red, and k = 4 is dark red. Vertical dotted lines indicate the k average nearest neighbor (kANN). (D) Pair correlation function computed as the average number of cells found between r and r + dr in comparison to a random point pattern (g > 1 indicates clustering, g < 1 indicates dispersion, and g ≈ 1 indicates randomness).

In the Euclidean space, the spatial distribution of stomata was largely preserved across all mutants. The NN distributions exhibited a peak at around 45, µm and the ANN and k-ANN were not significantly different from wild type (Fig. 3B, D and Fig. S2A, C). However, some subtle differences could be observed: the first peak at around 45, µm was more pronounced in *dek1-4* and *lgo-2* and the second peak at around 90, µm more pronounced in *lgo-2* (Fig. 3B), suggesting a slightly more regular local spacing in these mutants.

In the network space, however, more differences were observed, particularly in *dek1-4* and *lgo-2*. As the k-nearest neighbor distributions were shifted to the right, with a greater proportion of nearest stomata spaced by two cells and significantly higher ANN and k-ANN (Fig. 4B-C and Fig. S2B-D) in *dek1-4* and *lgo-2*, more pavement cells had formed between stomata complexes. The effect was more pronounced in *dek1-4* than in *lgo-2* despite *lgo-2* having more total cells. This could be due to the fact that newly formed cells in *lgo-2* can still enter the stomatal lineage [22], reducing the number of intercalated pavement cells between stomatal complexes.

In conclusion, reducing giant cell number results in more cells being intercalated between stomatal complexes in the tissue network, with only subtle effects on the spatial distribution of stomata within space. This indicates that stomatal spatial distribution is robust to such perturbation, which is consistent with stomatal number conservation. Whether the extra divisions occur before or after stomatal patterning remains unclear: if after, the robustness would reflect that these processes are temporally separated; if during, it would suggest that the stomatal patterning is unaffected by an increased number of available cells.

### Overexpression of *LGO* and *ATML1* affects the spatial pattern of stomata

In contrast to loss-of-function mutants, overexpression of *LGO* and *ATML1* significantly affects the stomatal spatial distribution, consistent with the observed reduction in stomatal number and density. In *LGO-OX*, stomata were more dispersed from each other, with higher k-ANN values and more spread k-ANN distributions (Fig. 3B-D and Fig. S2A, C), and more dispersed patterns in the pair correlation function at larger distances (g < 1). Locally, however, the first NN distribution still showed a peak at around 50, µm (Fig. 3B). Conversely, in *ATML1-OX*, stomata were more clustered, with lower k-ANN values but also a more heterogeneous pattern across the tissue, with peaks in the pair correlation function (g > 1) at about 40, µm, 110, µm, and 160, µm (Fig. 3C–D).

In the network space, both *LGO-OX* and *ATML1-OX* exhibited a reduced ANN (Fig. 4B and Fig. S2B). In *LGO-OX*, a greater proportion of nearest stomata were found one cell closer than in wild type (Fig. 4C), consistent with our previous observation that SLGCs can become giant cells in this genotype. If SLGCs endoreduplicate instead of completing spacing divisions, fewer cells would be expected between stomata. This suggests that the increased Euclidean spacing in *LGO-OX* is due to larger giant cell size but not to more intercalated cells. In *ATML1-OX*, stomata were observed more clustered in the network compared to wild type (Fig. 4C). This raises question on wether most non-giant cells are part of the stomatal lineage in this genotype.

In conclusion, both overexpression lines reduce stomatal number, but they affect stomatal spatial organization differently and potentially through distinct mechanisms: *LGO-OX* spread stomata through larger giant cell physical sizes, while *ATML1-OX* creates heterogeneous regions dense in stomata. These distinct spatial organizations, which likely have distinct consequences for stomatal function, were dissociable by using both network and point pattern analyses. This highlights the importance of considering the tissue context to understand how stomatal distribution is established.

### Giant cells reduce network path length and increase tissue network connectivity

Beyond the spatial pattern of stomata, we wondered whether the whole tissue exhibits distinct organizational features depending on the presence of giant cells. Giant cells are few but have a large number of neighbors. Therefore, we asked whether they might influence the global connectivity of the tissue network. Recently, network theory [24, 25, 26] has been used to describe global properties of cellular tissue organization, with some tissues exhibiting small-world network properties characterized by potential high transport efficiency [27, 28]. We used analogous observables [29] to investigate the global connectivity of the epidermal tissue network (Fig. 5).

**Figure 5:**
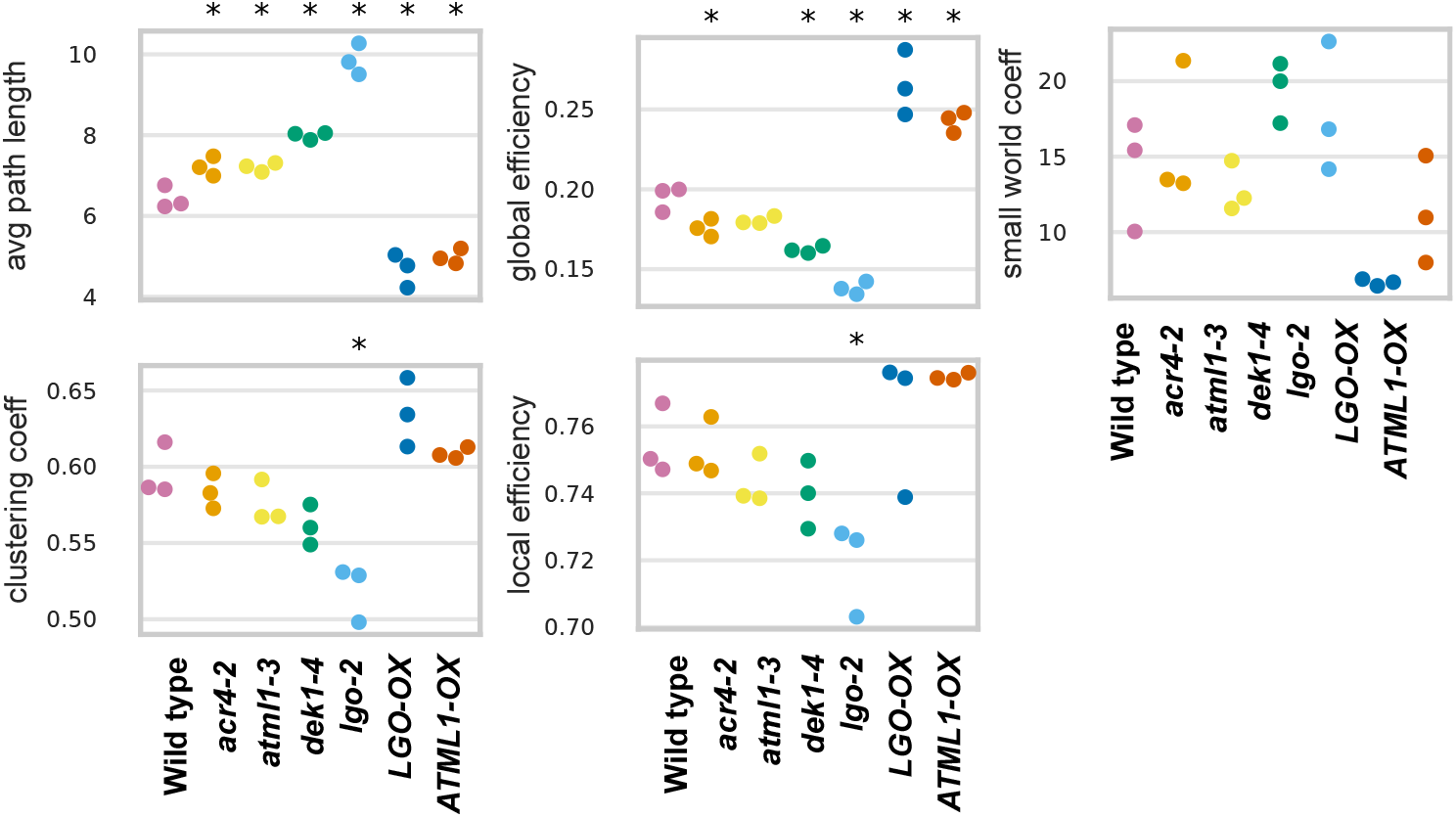
Giant cells reduce network path length and increase tissue network connectivity. Tissue level observables quantifying the connectivity between all cells in the tissue network [29] of the genotypes. (Left) Average path length between cells and clustering coefficient. (Middle) Global efficiency and local efficiency. (Right) Small world coefficient. Associated with Fig. 4.

We found that mutants lacking giant cells exhibited a significantly higher average path length between cells, such that the more small cells there are, the longer the average path between cells (Fig. 5). By contrast, the average path length was significantly reduced in *ATML1-OX* and *LGO-OX* compared with wild type, and to a similar extent. The clustering coefficient, reflecting the average number of neighbors per cell, followed the same trend, with mutants exhibiting fewer neighbors and overexpression lines more (Fig. 5). Accordingly, the global efficiency, which measures the extent to which cells are connected across the tissue [29], was significantly decreased in mutants and increased in overexpression lines (about 0.25 in *ATML1-OX* and 0.27 in *LGO-OX* compared with about 0.20 in wild type) (Fig. 5).

Collectively, these results show that giant cells, because of their large number of neighbors, act as hubs within the tissue network, reducing the path length between cells and in particular between stomatal complexes, which are otherwise relatively isolated from one another in wild-type. Whether this has functional consequences for cell-to-cell signaling remains unknown, but it suggests that the cell size heterogeneity produced by giant cells has tissue-level organizational consequences beyond local patterning.

## Discussion

We investigated how giant cell patterning affects stomatal number and spatial organization in the leaf epidermis. Our results show that stomatal number and density are robust to reduced numbers of giant cells but sensitive when giant cell formation is enhanced. Furthermore, in addition to the stomatal signaling pathway, the spatial distribution of stomata is also shaped by cell growth, cell division, and giant cell patterning, illustrating the interdependence between patterning systems in organizing the tissue.

We have shown quantitatively that the number of cells and tissue composition in the leaf epidermis is affected by the giant cell pathway. Overexpression of *ATML1* and *LGO* results in the production of fewer stomata and pavement cells, suggesting that entry into endoreduplication can compete for fate by preventing entry into the stomatal lineage. In addition, our spatial quantification suggests that *LGO* can promote giant cell formation in stomatal lineage cells. This is consistent with the previous finding [22] that *LGO* can promote endoreduplication in meristematic mother cells, possibly by reducing SPCH activity, but also in SLGC cells. In *ATML1-OX*, however, our spatial quantification suggests that stomatal entry is partially prevented by the giant cell pathway only in protodermal cells, but not in stomatal lineage cells.

In mutants lacking giant cells, the cell type composition was altered by an increase in the number of pavement cells, but the number of stomata was nearly the same as in wild type. This suggests that reduced endoreduplication favors cell division, resulting in more pavement cells, but does not necessarily promote stomatal entry. The robustness of stomatal number to changes in the total cell number in the epidermis could be consistent with a model in which SPCH functions in a continuous manner in space throughout the tissue, such that the number of available protodermal cells does not determine the number of stomatal entries, but rather the overall tissue area. Alternatively, timing of differentiation might be important. If divisions occur after stomatal entries, the robustness could simply reflect the sequential nature of these processes. Distinguishing between these scenarios would require live imaging of stomatal patterning dynamics in these mutants. It has been observed that the number of stomata in *lgo-2* is lower than in wild type at the beginning of tissue growth [22], whereas it is similar in the mature tissue [22], showing that newly formed cells can still enter the stomatal lineage. We also observed that despite having more cells in *lgo-2* compared to *dek1-4* or *atml1-3, lgo-2* had slightly fewer cells between stomata, which could be attributed to a decrease in the number of spacing and amplifying divisions of meristemoids.

By studying the stomatal spatial distribution and cellular arrangement across wild-type leaves, sepals, and genotypes involved in giant cell formation, we found that the stomatal pattern can be affected in multiple ways by giant cell patterning. Cell size and shape contribute to stomatal spacing independently of the patterning mechanism itself, as shown by the comparison between leaves and sepals, but also by the *LGO-OX* phenotype, where larger giant cells increase stomatal spacing in Euclidean space without altering their cellular arrangement in the network. By contrast, cell divisions regulated by the giant cell pathway alter the cellular arrangement in the network without disrupting the spatial distribution. Whether this reflects an active interaction with the stomatal pathway or a passive consequence of timing of differentiation is unclear.

Previous studies have often analyzed stomatal patterns from a point-pattern perspective alone, making it challenging to determine whether spatial differences result from cell shape or topological arrangement. Our results show that analyzing both the spatial distribution in the Euclidean space and the cellular arrangement in the network space provides complementary and necessary information to understand cellular patterning. Collectively, our results indicate that both cell division and cell growth, including giant cell formation, shape the stomatal spatial pattern. This likely has consequences for stomatal function, as stomatal density, spacing, and distribution are known to influence gas exchange, water use efficiency, and guard cell mechanics [10, 11, 12]. Stomatal patterning and giant cell patterning mutually influence each other: proliferative cells have also been shown to shape the giant cell pattern over time [5]. Therefore, in tissues composed of multiple cell types, different patterning systems are interconnected and influence each other during tissue growth. This highlights the importance of considering the broader tissue context and the interplay between patterning systems to fully understand how cellular organization is established.

## Methods

### Plant images

We reused the datasets from [5] for the new analyses performed in this manuscript.

### Image processing and analysis

All the image segmentation and cell type classification were performed using the same datasets and analysis pipelines as described in [5]. No additional image acquisition or reprocessing was performed.

### Cell type composition analysis

The tissue cell type composition in each experimental replicate was analyzed by computing the number and fraction of each cell type. The fraction was calculated within the segmented regions of the leaf (Fig. 1), and within a selected rectangular region of the sepal (Fig. 1) to exclude cells at the edge and bottom of the sepal, which lack stomata. This approach ensured a fair comparison between the two organs. The fraction for each cell type ct (Fig. 1) was calculated as 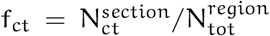, where N_ct_ is the number of cells of type ct in the segmented region, and N_tot_ is the total number of cells within the region. The number of cells of type ct in the entire abaxial epidermis was computed using the entire segmented epidermis in the sepal. In the leaf, as it was not possible to obtain images of the entire leaf at 25 dpg, the total number of cells of type ct 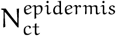 in the abaxial epidermis was estimated in each replicate by using the leaf size as follows: 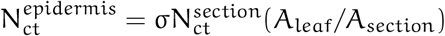, where A_leaf_ is the area of the leaf, A_section_ is the area of the segmented cells, and σis the fraction of the leaf occupied by the leaf blade, which we estimated to be σ ≈ 0.9.

### Quantification of spatial cellular patterns

The quantification of the cellular spatial patterns were conducted using two distinct perspectives: the point pattern point of view, using cell center coordinates, and the network point of view, where nodes represent cells and edges represent intercellular connections [30]. The cell coordinates and cell graphs were computed using MorphoGraphX [31, 32]. A custom Python script was employed to perform all the analyses.

The pairwise d_ij_ distances between all cells of a given cell type ct (in this case, stomata) were calculated. The distances were calculated in both the Euclidean space, by computing the Euclidean distance between point coordinates, and in the network space, by computing the shortest path in number of edges between cells by using the NetworkX package [33]. For each cell i, the nearest neighbor distance was computed as

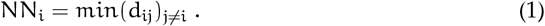

To quantify the spatial distributions of the cell population locally, the distribution of the nearest neighbor distances 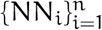 was studied, and the average nearest neighbor distance ANN was then calculated as

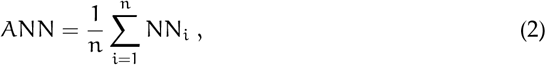

with n the number of cells of type ct. The k-nearest neighbor kNN_i_ for each cell i was also computed for k = 2, 3 and 4, as the k-th smallest distance in 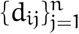, and the average k nearest neighbor distance kANN was calculated. To quantify the variability of the cell local spatial distributions, the coefficient of variation (CV) was calculated for the first NN as 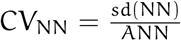, where sd(NN) is the standard deviation of the distribution and ANN is the average.

To study the spatial organization of cells at a higher level, the average number of cells at a distance r (or d in the network space, with d representing the number of edges) was calculated as follows:

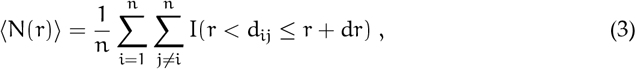

where I(r < d_ij_ ≤ r + dr) is an indicator function equal to 1 when the distance between a pair of cells d_ij_ lies between r and r + dr, and 0 otherwise. This was compared to the average number of cells observed between a distance r and r + dr in a random reference, denoted ⟨N_random_(r)⟩, which is the mean of the average number of cells at distance r across multiple random simulations, to build an pair correlation function g as

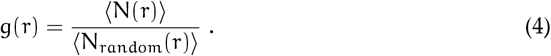

Similarly to a radial distribution function, the pair correlation function g is used to quantify the correlation between all the pairs of cells, indicating whether cells are more clustered (g > 1) or less densely distributed (g < 1) compared to what is expected randomly within each distance interval r and r + dr.

To statistically test the randomness of the stomatal patterns in the wild-type leaves, the same method as in [5] was used, by using the randomized tissues as a null model (Fig. **??**). N_random_(r) was calculated in every randomized tissue, and the pair correlation function function was used as defined in Eq. 4.

In order to compare the stomatal patterns between leaves and sepals (Fig. 2) and between genotypes (Fig. 3 and Fig. 4), the null model were estimated without the help of the randomized tissues. For point pattern analysis, N_random_ was obtained by using a random point pattern as a null model, utilizing the Python package pointpats [34]. For the network analysis, N_random_ was estimated by computing the average number of cells at each edge distance of a stomata. This was followed by the calculation of the expected number of stomata at each distance d as if they were placed randomly on the graph. In addition, all of the observables (the NN distribution, the kNN distributions, the ANN, the kANN, and the pair correlation function g) were computed in individual replicates (Fig. S2) and in the pooled replicates (were all distances were considered together) (Fig. 3 and Fig. 4).

In addition to the distance-based measures, several metrics derived from network theory [29, 24, 26] were calculated and averaged over all cells by using the Python package NetworkX. These metrics were computed in different genotypes to assess the global efficiency of the cellular connectivity (Fig. 5). The average path length is defined as the average path length between all pair of cells. Each path length is the shortest path between two cells. The clustering coefficient is computed as the average number of neighbors (or degree) in the graph. The global efficiency, measuring the efficiency of communication across the whole cellular network, is defined as the average of inverse distances (shortest path length) between all pairs of nodes in the network. The local efficiency, measuring the robustness to random failure of nodes, is the global efficiency computed in the neighboring nodes of each nodes and averaged over all nodes. The small-world coefficient is defined as the ratio of the clustering coefficient of the network to the clustering coefficient of a random network, normalized by the ratio of the average path length of the network to the one of a random network. A high small-world coefficient indicates both high clustering and short average path lengths.

## Data availability

The confocal images analysed were generated from the study in [5], with a few two leaf replicates for *lgo-2* and *LGO-OX* and two wild-types, which are datasets from [35]. As reported in [5], the microscopy data, MorphoGraphX meshes of the images used in this manuscript, and cell classification and randomization are available at Open Science Framework (osf.io), DOI: https://doi.org/10.17605/OSF.IO/RFCWS. Code for the new data analyses performed in this manuscript to reproduce the figure results can be provided upon request.

## Author contributions

GW and PF-J conceived the work. GW performed the analyses shown in the manuscript. GW, FKC, AHKR, PF-J interpreted the results. GW wrote the first draft. GW, FKC, AHKR, PF-J revised and edited the manuscript. PF-J supervised the work.

## Declaration of interests

The authors declare no competing interests.

## Acknowledgements

We would like to acknowledge John Chandler for critical comments on the manuscript, and to Tobias Bollenbach, Aneta Koseska and Michael Raissig for fruitful discussions. This work was funded by the International Max Planck Research School (https://www.mpipz.mpg.de/imprs/) on “Understanding Complex Plant Traits Using Computational and Evolutionary Approaches” (GW); a core grant from the Max Planck Society (https://www.mpg.de/) (PF-J); the Deutsche Forschungsgemeinschaft (DFG, German Research Foundation, https://www.dfg.de) under Germany’s Excellence Strategy— EXC-2048/1—project ID 390686111 (PF-J); and the National Science Foundation (US NSF, https://www.nsf.gov), the Division of Biological Infrastructure DBI-232051 (PF-J, AHKR).

## Notes

### Competing Interest Statement

The authors have declared no competing interest.

## References

[1] Bergmann DC, Sack FD. Stomatal development. Annual Review of Plant Biology. 2007;58:163–81.

[2] Torii KU. Two-dimensional spatial patterning in developmental systems. Trends in cell biology. 2012;22(8):438–46.

[3] Roeder AHK, Chickarmane V, Cunha A, Obara B, Manjunath BS, Meyerowitz EM. Variability in the Control of Cell Division Underlies Sepal Epidermal Patterning in Arabidopsis thaliana. PLOS Biology. 2010;8(5):e1000367. Available from: https://journals.plos.org/plosbiology/article?id=10.1371/journal.pbio.1000367.

[4] Roeder AH, Cunha A, Ohno CK, Meyerowitz EM. Cell cycle regulates cell type in the Arabidopsis sepal. Development. 2012;139(23):4416–27.

[5] Clark FK, Weissbart G, Wang X, Harline K, Li CB, Formosa-Jordan P, et al. A common pathway controls cell size in the sepal and leaf epidermis leading to a nonrandom pattern of giant cells. PLOS Biology. 2025;23(11):e3003469.

[6] Zuch DT, Doyle SM, Majda M, Smith RS, Robert S, Torii KU. Cell biology of the leaf epidermis: Fate specification, morphogenesis, and coordination. The Plant Cell. 2022;34(1):209–27.

[7] Dong J, Bergmann DC. Stomatal patterning and development. Current topics in developmental biology. 2010;91:267–97.

[8] Torii KU. Stomatal development in the context of epidermal tissues. Annals of botany. 2021 jul;128(2):137–48. Available from: https://pubmed.ncbi.nlm.nih.gov/33877316/.

[9] Robinson DO, Coate JE, Singh A, Hong L, Bush M, Doyle JJ, et al. Ploidy and Size at Multiple Scales in the Arabidopsis Sepal. The Plant Cell. 2018 nov;30(10):2308–29. Available from: https://dx.doi.org/10.1105/tpc.18.00344.

[10] Dunn J, Hunt L, Afsharinafar M, Meselmani MA, Mitchell A, Howells R, et al. Reduced stomatal density in bread wheat leads to increased water-use efficiency. Journal of Experimental Botany. 2019 Jun;70(18):4737–4748. Available from: http://dx.doi.org/10.1093/jxb/erz248.

[11] Harrison EL, Arce Cubas L, Gray JE, Hepworth C. The influence of stomatal morphology and distribution on photosynthetic gas exchange. The Plant Journal. 2019 Nov;101(4):768–779. Available from: http://dx.doi.org/10.1111/tpj.14560.

[12] Papanatsiou M, Amtmann A, Blatt MR. Stomatal Spacing Safeguards Stomatal Dynamics by Facilitating Guard Cell Ion Transport Independent of the Epidermal Solute Reservoir. Plant Physiology. 2016 Jul;172(1):254–263. Available from: http://dx.doi.org/10.1104/pp.16.00850.

[13] Trozzi N, Majda M. How stochastic cell fate and endoreduplication yield nonrandom epidermal patterns. Quantitative Plant Biology. 2026;4:1–4.

[14] Pyke K, Marrison J, Leech A. Temporal and spatial development of the cells of the expanding first leaf of Arabidopsis thaliana (L.) Heynh. Journal of experimental botany. 1991;42(11):1407–16.

[15] Ezaki K, Koga H, Takeda-Kamiya N, Toyooka K, Higaki T, Sakamoto S, et al. Precocious cell differentiation occurs in proliferating cells in leaf primordia in Arabidopsis angustifolia3 mutant. Frontiers in Plant Science. 2024;15:1322223.

[16] Donnelly PM, Bonetta D, Tsukaya H, Dengler RE, Dengler NG. Cell Cycling and Cell Enlargement in Developing Leaves of Arabidopsis. Developmental Biology. 1999 nov;215(2):407–19.

[17] Andriankaja M, Dhondt S, De Bodt S, Vanhaeren H, Coppens F, De Milde L, et al. Exit from proliferation during leaf development in Arabidopsis thaliana: a not-so-gradual process. Developmental cell. 2012;22(1):64–78.

[18] Geisler M, Nadeau J, Sack FD. Oriented Asymmetric Divisions That Generate the Stomatal Spacing Pattern in Arabidopsis Are Disrupted by the too many mouths Mutation. The Plant Cell. 2000 Nov;12(11):2075–2086. Available from: http://dx.doi.org/10.1105/tpc.12.11.2075.

[19] Shi P, Jiao Y, Diggle PJ, Turner R, Wang R, Niinemets Ü. Spatial distribution characteristics of stomata at the areole level in Michelia cavaleriei var. platypetala (Magnoliaceae). Annals of Botany. 2021 nov;128(7):875–86. Available from: https://dx.doi.org/10.1093/aob/mcab106.

[20] Liu C, Li Y, Xu L, Li M, Wang J, Yan P, et al. Stomatal Arrangement Pattern: A New Direction to Explore Plant Adaptation and Evolution. Frontiers in Plant Science. 2021;Volume 12 - 2021. Available from: https://www.frontiersin.org/journals/plant-science/articles/10.3389/fpls.2021.655255.

[21] Tong Yang K, peng Chen G, ren Xian J. Stomatal distribution pattern for 90 species in Loess Plateau – Based on replicated spatial analysis. Ecological Indicators. 2023;148:110120. Available from: https://www.sciencedirect.com/science/article/pii/S1470160X23002625.

[22] Dubois M, Achon I, Brench RA, Polyn S, Tenorio Berrío R, Vercauteren I, et al. SIAMESE-RELATED1 imposes differentiation of stomatal lineage ground cells into pavement cells. Nature Plants. 2023;9(7):1143–53.

[23] Laruelle E, Spassky N, Genovesio A. Unraveling spatial cellular pattern by computational tissue shuffling. Communications Biology. 2020;3(1):605.

[24] Barabási AL, Oltvai ZN. Network biology: Understanding the cell’s functional organization; 2004.

[25] Bullmore E, Sporns O. Complex brain networks: graph theoretical analysis of structural and functional systems. Nature Reviews Neuroscience. 2009 Feb;10(3):186–198. Available from: http://dx.doi.org/10.1038/nrn2575.

[26] Strogatz SH. Exploring complex networks. Nature. 2001 Mar;410(6825):268–276. Available from: http://dx.doi.org/10.1038/35065725.

[27] Novkovic M, Onder L, Bocharov G, Ludewig B. Topological Structure and Robustness of the Lymph Node Conduit System. Cell Reports. 2020 Jan;30(3):893–904.e6. Available from: http://dx.doi.org/10.1016/j.celrep.2019.12.070.

[28] Lee MD, Buckley C, Zhang X, Louhivuori L, Uhlén P, Wilson C, et al. Small-world connectivity dictates collective endothelial cell signaling. Proceedings of the National Academy of Sciences. 2022 Apr;119(18). Available from: http://dx.doi.org/10.1073/pnas.2118927119.

[29] Bassel GW. Multicellular Systems Biology: Quantifying Cellular Patterning and Function in Plant Organs Using Network Science. Molecular Plant. 2019;12(6):731–42. Available from: 10.1016/j.molp.2019.02.004.

[30] Weissbart G, Formosa-Jordan P. Quantifying cellular spatial organization to infer biological mechanisms; 2026. HAL Preprint. Available from: https://hal.science/hal-05602235.

[31] Barbier de Reuille P, Routier-Kierzkowska AL, Kierzkowski D, Bassel GW, Schüpbach T, Tauriello G, et al. MorphoGraphX: A platform for quantifying morphogenesis in 4D. elife. 2015;4:e05864.

[32] Strauss S, Runions A, Lane B, Eschweiler D, Bajpai N, Trozzi N, et al. Using positional information to provide context for biological image analysis with MorphoGraphX 2.0. Elife. 2022;11:e72601.

[33] Hagberg AA, Schult DA, Swart PJ. Exploring Network Structure, Dynamics, and Function using NetworkX. In: Varoquaux G, Vaught T, Millman J, editors. Proceedings of the 7th Python in Science Conference. Pasadena, CA USA; 2008. p. 11 15.

[34] Kang W, Wolf LJ, Rey S, Shao H, Mridul Seth, Fleischmann M, et al. pysal/pointpats: pointpats 2.3.0. Zenodo; 2023. Available from: https://zenodo.org/record/7706219.

[35] Trozzi N, Lane B, Perruchoud A, Wang Y, Hoermayer L, Ansel M, et al. Puzzle cell shape emerges from the interaction of growth with mechanical constraints. bioRxiv. 2023.

