## Supplementary information for "Stomatal patterning is shaped by the interplay with giant cell patterning in Arabidopsis"

### Supplementary figures

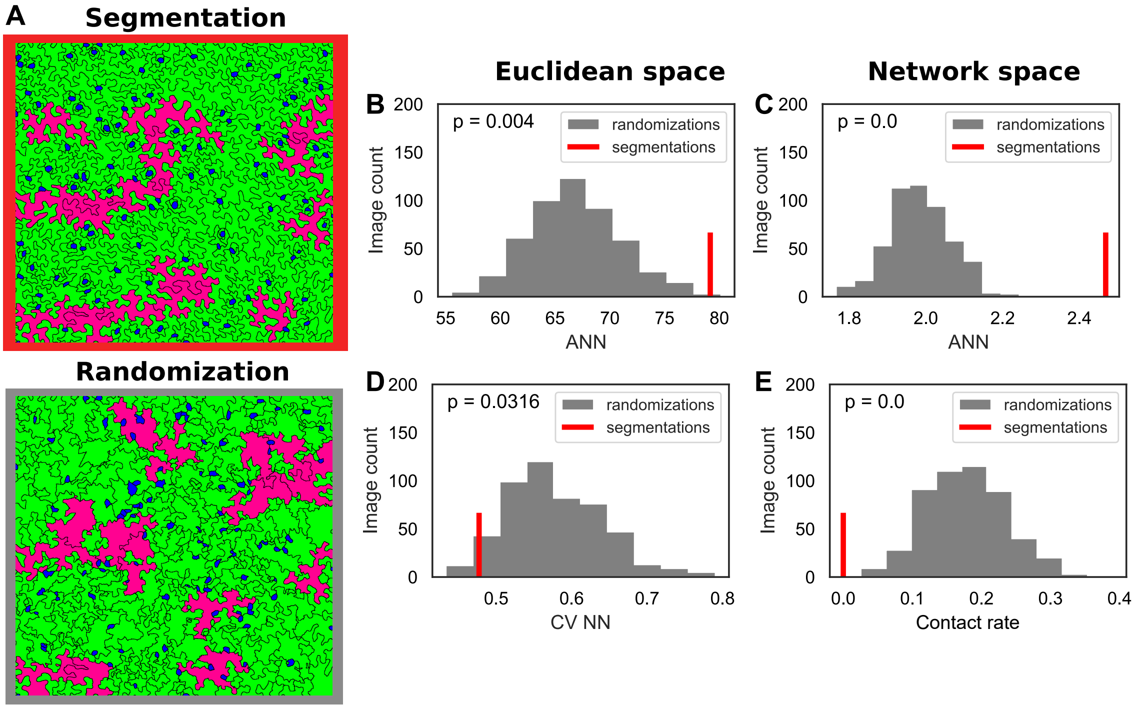


Fig. S1: **Statistical analysis of the randomness of the local stomatal pattern in wild-type.** (A) Example of a representative segmentation of a wild-type leaf 25 dpg (segmentation) and one corresponding randomized tissue image (randomization). To statistically assess the extent to which the segmented image shows a random stomatal pattern, a quantitative observable was extracted from the segmentation and compared with the same observable computed in all randomized tissues. (B–C) Average nearest neighbor (ANN) distance in the Euclidean space (B) and in the network space (C). (D) Coefficient of variation of the nearest neighbor distance (CV NN) in the Euclidean space. (E) Contact rate between stomata in the cellular tissue. The mean value observed in the segmentation (represented by the red bar) is significantly different from the values expected by chance (indicated by the grey distribution) in random tissues in (B–D).

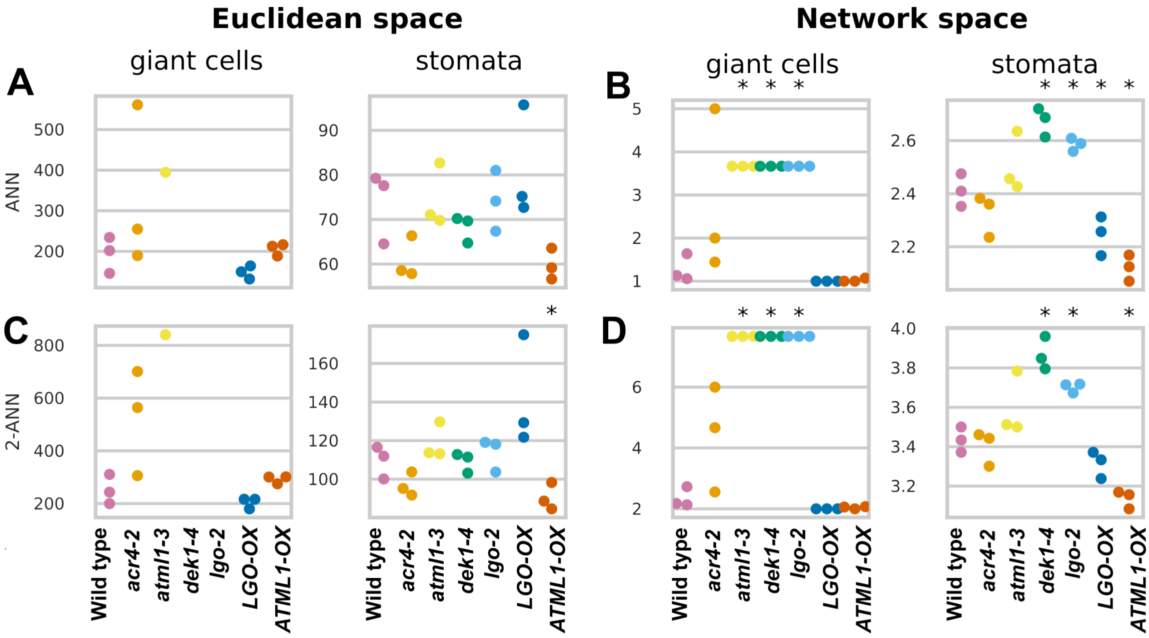


Fig. S2: **Mean nearest neighbor observables of stomata and giant cells in genotypes.** (A–B) First average nearest neighbor (ANN) distance in the Euclidean space (A) and in the network space (B). (C–D) Second ANN in the Euclidean space (C) and in the network space (D).
